# Checkpoint non-fidelity induces a complex landscape of lineage fitness after DNA damage

**DOI:** 10.1101/431486

**Authors:** Callum J. Campbell, Ashok R. Venkitaraman, Alessandro Esposito

## Abstract

DNA damage in proliferating mammalian cells causes death^1^, senescence^2^ or continued survival, via checkpoints that monitor damage and regulate cell cycle progression, DNA repair and fate determination^3^. Cell cycle checkpoints facilitate tumour suppression by preventing the generation of proliferating mutated cells^4^, particularly by blocking passage of DNA lesions into replication and mitosis^5^. While checkpoint non-fidelity permits cells to carry genomic aberrations into subsequent cell cycle phases^6^, its long-term consequences on lineages descendant from damaged cells remains poorly characterised. Devising methods for microscopy-based lineage tracing, we unexpectedly demonstrate that transient DNA damage to single living cells bearing a negligent checkpoint induces heterogenous cell-fate outcomes in their descendant generations removed from the initial insult. After transiently damaged cells undergo an initial arrest, pairs of descendant cells without obvious cell-cycle abnormalities either divide or die in a seemingly stochastic way. Progeny of transiently damaged cells may die generations afterwards, creating considerable variability of lineage fitness that promotes overall persistence in a mutagenic environment. Descendants of damaged cells frequently form micronuclei, activating immunogenic signalling. Our findings reveal previously unrecognized, heterogenous effects of cellular DNA damage that manifest long afterwards in descendant cells. We suggest that these heterogenous descendant cell-fate responses may function physiologically to ensure the elimination and immune clearance of damaged cell lineages, but pathologically, may enable the prolonged survival of cells bearing mutagenic damage.

## Introduction

DNA damage in mammalian cells triggers many co-ordinated processes collectively known as the DNA damage response (DDR). These allow cell cycle checkpoints to delay cell cycle progression permitting the intervention of DNA repair and cell fate determination mechanisms, so facilitating the maintenance of genomic integrity^4^. Despite this, several studies have suggested that both non-transformed and cancer cells exhibit varying degree of ‘checkpoint negligence’ dependent on the timing and the amount of DNA damage^6–8^. For instance, the G2 checkpoint is surprisingly lax in preventing cells entering mitosis with DNA damage^9^ while the G1 checkpoint becomes relatively insensitive to activation the later in G1 DNA damage occurs^10–13^. It is also now apparent that the G2 checkpoint might not be directly controlled by a threshold of unrepaired DNA damage. Instead, mitotic entry appears to be controlled by the accumulation of substrates phosphorylated by Polo-like kinase 1 (PLK1)^14,15^. Although the rate of accumulation is controlled by the DDR, heterogeneity in PLK1 activity results in significant cell-to-cell variability in the fidelity and timing of entry into mitosis. In contrast with the detailed knowledge we have on molecular mechanisms underlying checkpoints, the characterisation of the extent and the consequences of descendant cells inheriting unrepaired lesions remains remarkably limited. Such characterisation is necessary to reconcile the apparent fallibility of cell cycle checkpoints with the crucial role in maintaining genomic integrity revealed by loss of function studies. Crucially, a better understanding of cell fate determination during the DNA damage response is of significant therapeutic importance given the use of DNA damage in chemotherapies, where efficacy is often limited by fractional killing and the emergence of resistant phenotypes^16^.

Aiming to distinguish between different cell fates and assess the role of checkpoint cooperation and non-genetic heterogeneity in checkpoint fidelity, we developed single-cell microscopy-based lineage tracing of DNA damaged U2OS cells, a well-characterised cancer cell line. While addressing a simple but fundamental question about what happens to the descendants of DNA damaged cells that progress through the cell cycle, we reveal a more complex response to DNA damage than previously described. After an initial cell cycle arrest, cells continue cycling with DNA damage, often without showing abnormalities in cell cycle timing, but do trigger tumour suppressive mechanisms in later generations. We offer a fresh and provocative perspective whereby the non-genetic heterogeneity in checkpoint fidelity previously identified in U2OS cells^14,15^ causes a profound heterogeneity of lineage fitness in response to DNA damage; this heterogeneity, invisible to population measurements and revealed by single-cell lineage tracing, poses challenges in the design of therapeutic interventions, but also opportunities.

## Results

### Lineage tracing of U2OS cells after DNA damage

To quantify the effects of DNA damage on cells progressing through checkpoints with unresolved lesions, we engineered a U2OS cell line stably expressing the FUCCI probes^17^ and a nuclear marker (U2OS-Tracer) (**Fig. 1**, **Extended Data Fig. 1 and Supp. Methods**). The FUCCI probes permitted us to assess the phase of the cell-cycle when each cell experienced DNA damage, to quantify the consequent effects on the length of G1 or S/G2/M in the treated cell and in its descended lineage. Lineage tracing and correlation analysis required the *ad hoc* development of plasmids and analytical tools (LinSense and CloE-DDR), shared in public repositories (see **Methods**), for the tracking of cell lineages, cell cycle determination, manual curation of lineages and modelling of clonal evolution.

**Figure 1.**
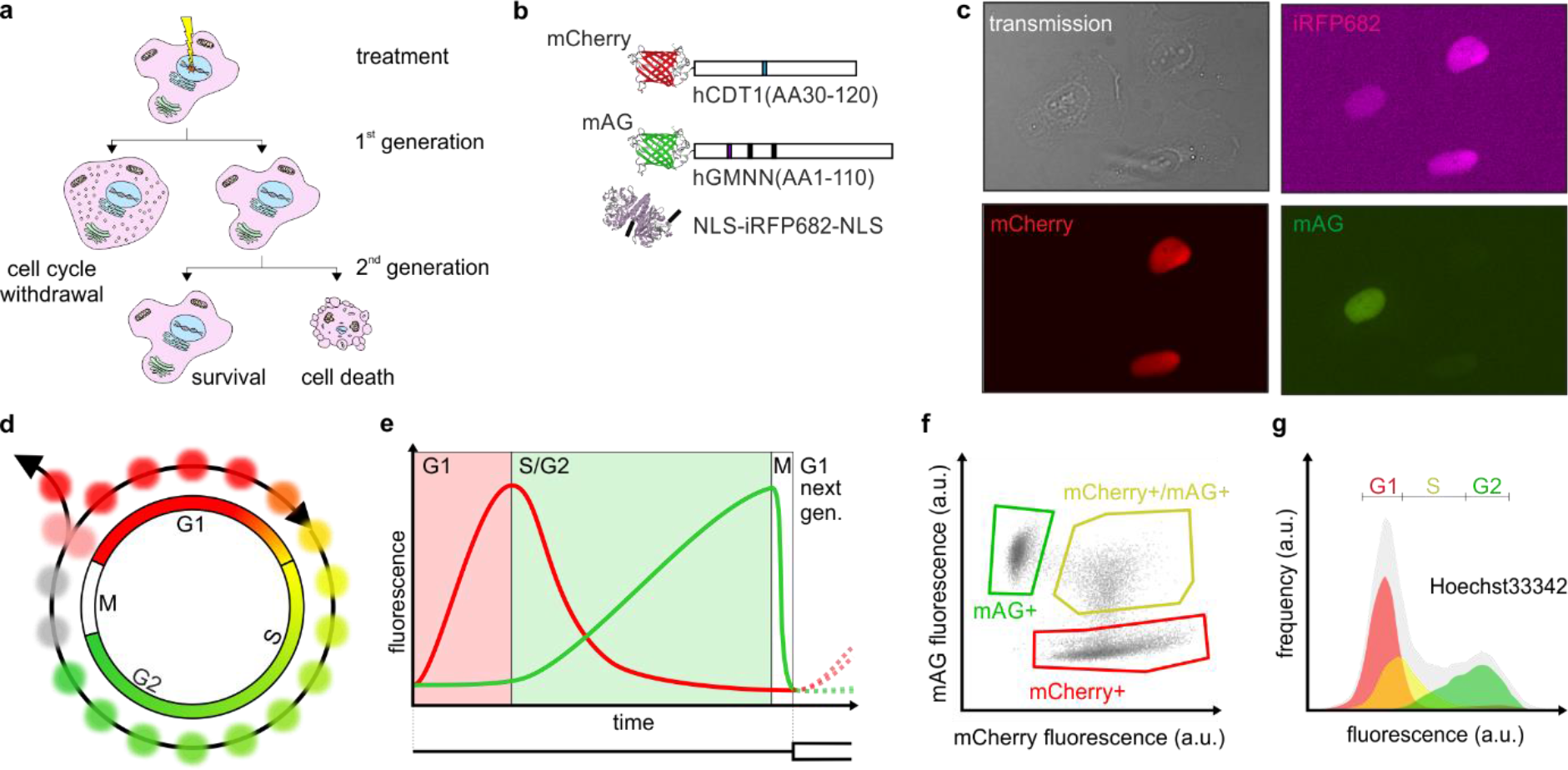
Lineage analysis of U2OS FUCCI-NM cells. To track cell fate choices (*e.g.*, arrest, proliferation, senescence or cell death) after DNA damage during the cell cycle (**a**), we engineered U2OS cells to stably express a FUCCI system made of mCherry-hCDT1(aa30-120), mAG-hGeminin(aa1-110) and NLS-IRFP682-NLS (**b**). U2OS FUCCI-NM cells constitutively express the fluorescent nuclear marker while the fluorescent red and green FUCCI probes (**c**) vary as a function of cell cycle (**d**). We used the peak of red fluorescence and transition to green fluorescence to mark the G1-S transition, while using the nuclear markers and transmission images to aid generational tracing (**e**). Gating green and red fluorescent cells (**f**) in flow cytometry, we could confirm with Hoecst33342 staining (**g**) that the U2OS-Tracer cells faithfully report cell cycle progression.

### DNA damage drives cell death in descendant U2OS cells

U2OS-Tracer cells were imaged for six days continuously with time-lapse microscopy and treated with the radiomimetic drug neocarzinostatin (NCS). NCS induces DNA damage in all treated cells (**Extended Data Fig. 2**) that are then imaged for a further 5 days. LinSense then permitted us to map cell fates (proliferation, cell death, survival), cell cycle lengths and relationships between cells, for hundreds of mock- and NCS-treated cell per experimental repeat (**Extended Data Fig. 3**). Hereafter, the treated cells, their daughters and granddaughters for mock and NCS treated lineages are referred to as: mock-treated, mock-daughters, mock-granddaughters, NCS-treated, NCS-daughters and NCS-granddaughters, respectively.

The cell cycle length of mock-treated cells (**Fig. 2**, 23±3hrs; median cell cycle length ± median absolute deviation), mock-daughters cells (20±3hrs) and mock-granddaughters cells (19±2hrs) are similar. All three mock generations show a similar small proportion of dying cells. Both observations are consistent with relatively unperturbed cells behaving similarly in each generation during long-term imaging.

**Figure 2.**
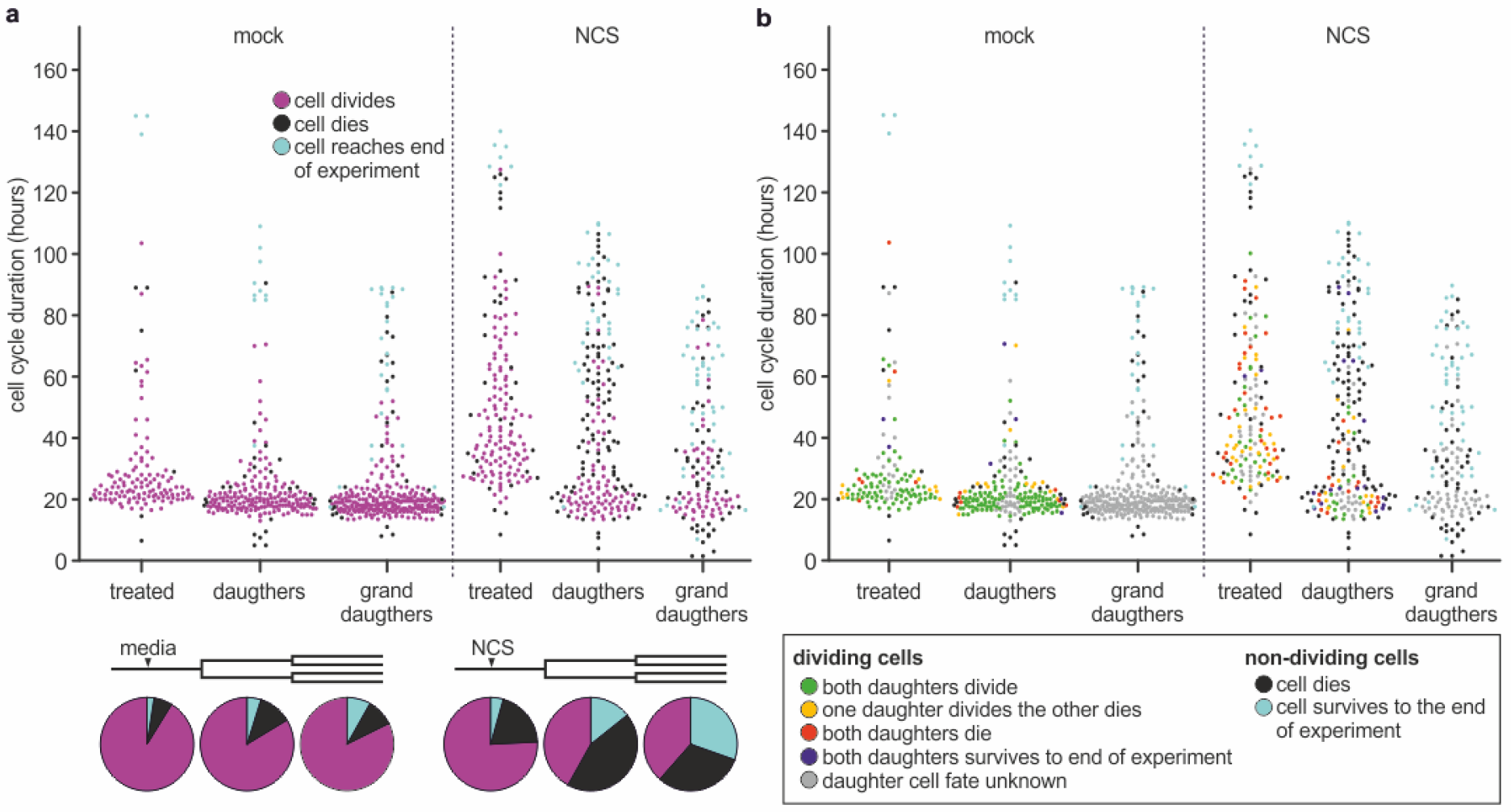
Generational analysis of DNA damaged cells. The total cell cycle length of mock- (**a**, left) and NCS- (**a**, right) are shown as a swarm plot where different colours identify cells that, from within each analysed generation, divide (magenta), die (black) or survive to the end of the experiment (5 days). The bottom pie charts show the prevalence of the three phenotypes for each generation. From the first generation after NCS treatment, cell cycle length for dividing cells converges to the normal ~20hr cell cycle for U2OS-Tracer cells. Recolouring these cells by the fate of their own daughters (**b**, legend inset) reveals the highly overlapping green, yellow and red populations in NCS-Daughters suggesting there are no obvious cell cycle alterations that predict the fate of the resultant NCS-Granddaughters.

As expected, NCS-treated cells exhibit a significant cell cycle arrest (**Fig. 2a** and **Extended Data Fig. 2**) resulting in much longer cell cycle durations (41±12hrs) and an increase in the proportion of cells that die (20% vs 6%) relative to NCS-mock cells. This observation confirms that U2OS cells retain a functioning DNA damage response. Nonetheless, U2OS are known to carry residual DNA damage through mitosis^14^ and we expected this to manifest as changes in the later generations. Indeed, we observed an increase in cell death proportions in the NCS-daughters (44% vs 12%) and NCS-granddaughters (31% vs 10%) generations relative to their mock counterparts, a phenomenon we termed ‘delayed cell death’. Furthermore, we observed that the duration of the initial cell cycle arrest appeared to correlate with the cell cycle timing of the treatment (**Fig. 3a-b**). The later in SG2M DNA damage occurs, the shorter the duration of the cell cycle arrest that can be induced, consistent with previous work indicating a progressive commitment to mitosis during G2 which results in non-genetic heterogeneity in the G2 checkpoint^14^.

**Figure 3.**
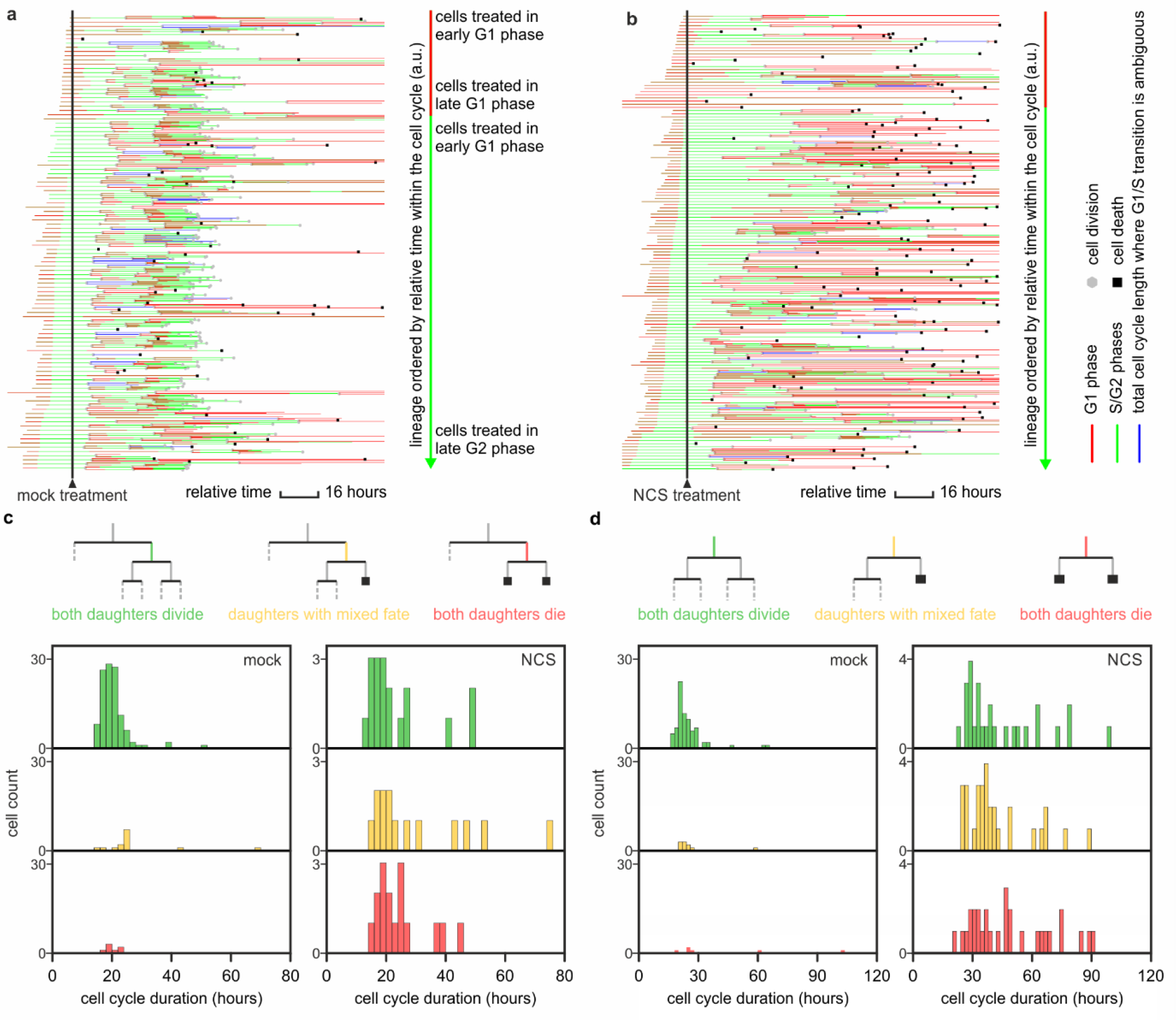
Cell-to-cell variability in the response to DNA damage. Cell-to-cell variability in the response to DNA damage Lineages of mock-treated cells exhibit moderate heterogeneity (**a**) but NCS-treated cells give raise to highly heterogeneous responses with what appears to be stochastic cell death at different generations (**b**) showing no correlation between cell cycle arrest and cell fate in across generations, either between daughter-granddaughters (**c**) or treated-daughter (**d**) cells.

### DNA damage causes considerable heterogeneity in cell proliferation and cell death

Following these observations, we hypothesised that this ‘delayed cell death’ phenotype was a mere consequence DNA damage inherited by permissive checkpoints that temporary arrest the cell cycle at each generation until the damage is either resolved or a cell dies. Consistent with this hypothesis, NCS-daughter cells that die, often do so following a prolonged G1 phase (**Extended Data Fig. 4**). This observation supports the notion that a permissive G2 DNA damage checkpoint can be compensated by cooperating with a more robust G1 checkpoint^6^.

Furthermore, we expected many of the dividing NCS-daughters cells to exhibit evidence of cell cycle arrests indicating the presence of damaged DNA that will then be inherited by NCS-granddaughters causing those to die. However, conflicting with this interpretation, the distribution of cell cycle length values of dividing NCS-daughters cells is not only significantly shorter than the arrested NCS-treated cells, but exhibits a distribution of cell cycle lengths that is predominantly normal as defined by their mock treated counterparts (**Fig. 2a**). We further analysed correlations between cell cycle lengths and cell fates, looking for evidence that delayed cell death is caused by inherited DNA damage and checkpoint cooperation. We expected the distribution of cell cycle length for dividing NCS-daughters cells that produce NCS-granddaughters cells that die to be significantly longer than dividing NCS-daughters that produce NCS-granddaughters that go on to divide. However, there is no apparent correlation between cell cycle length and cell fate in the subsequent generation (**Fig. 2b**); furthermore, the distribution of cell cycle lengths for those NCS-daughters cells with two daughter cells (NCS-granddaughters) that either both die (**Fig. 3c**, red), both survive (**Fig. 3c**, green) or exhibit mixed cell fates (**Fig. 3c**, yellow) are statistically identical (two-sample pairwise KS-test at 5% significance level with multiple comparison Bonferroni correction). Dying NCS-granddaughters are therefore frequently born of cells with no obvious abnormalities in cell cycle duration that would indicate inherited DNA damage activating cell cycle checkpoints (**Fig. 3d**).

To exclude that these observations are caused by the prolonged imaging of cells in NCS-treated conditions, we performed a surrogate lineage tracing experiment with parental U2OS cells and flow cytometry-based assay (**Extended Data Fig. 5**). Although flow cytometry does not permit measurement of all features of lineages traced by microscopy, this assay confirmed that parental U2OS cells, upon DNA damage, also resume cycling after an initial arrest while shedding a fraction of cells at each cell cycle (See **Supp. Note. 1**). This data suggest that DNA damage is propagated across cell generations causing death in the NCS-daughters and NCS-granddaughters, without triggering cell cycle arrest in the dividing NCS-daughters generation. Taken together, these observations suggest that in several cell lines, DNA damage alters cell fate over multiple generations, presumably due to inheritance and propagation of DNA damage itself or through more indirect means.

### DNA damage results in dramatically lowered but heterogeneous lineage fitness

The seemingly stochastic behaviour described in the generational analyses results in a considerable heterogeneity in cell proliferation and cell death between lineages (**Fig. 3a, b** and **Extended Data Fig. 3**). However, generational analysis alone fails to describe the consequences of this observation. Lineages vary in ‘fitness’, a concept which includes both how quickly cells proliferate and how many cells in a lineage survive. Here, we introduce a method for scoring lineage fitness utilizing a combination of two metrics, a proliferation index (PI, a weighted measure of how many generations a lineage passes through) and a survival index (SI, the fraction of the lineage that survives).

Examples of how different lineages are scored are given in **Extended Data Fig. 6** and **Supp. Note 2**. Mock treated cells typically give rise to lineages exhibiting high PI and SI scores (**Fig. 4a**), indicating the expected level of high fitness for relatively unperturbed cells. Predictably, NCS treatment results in large drops in both PI and SI scores indicating a significant effect on lineage fitness (**Fig. 4b**). However, the representation of fitness in a two-dimensional ‘lineage space’ permits to appreciate the consequences of a non-ideal checkpoint. Contrary to the simplistic depiction of fractional killing, where we classify cells in binary classes (cells die or survive), the fitness of lineages is very heterogeneous ranging from values where the founder cell itself dies to complete lineages and intermediate cases where a linage is terminally exhausted or support significant fitness by intermediate levels of proliferation and survivals.

**Figure 4.**
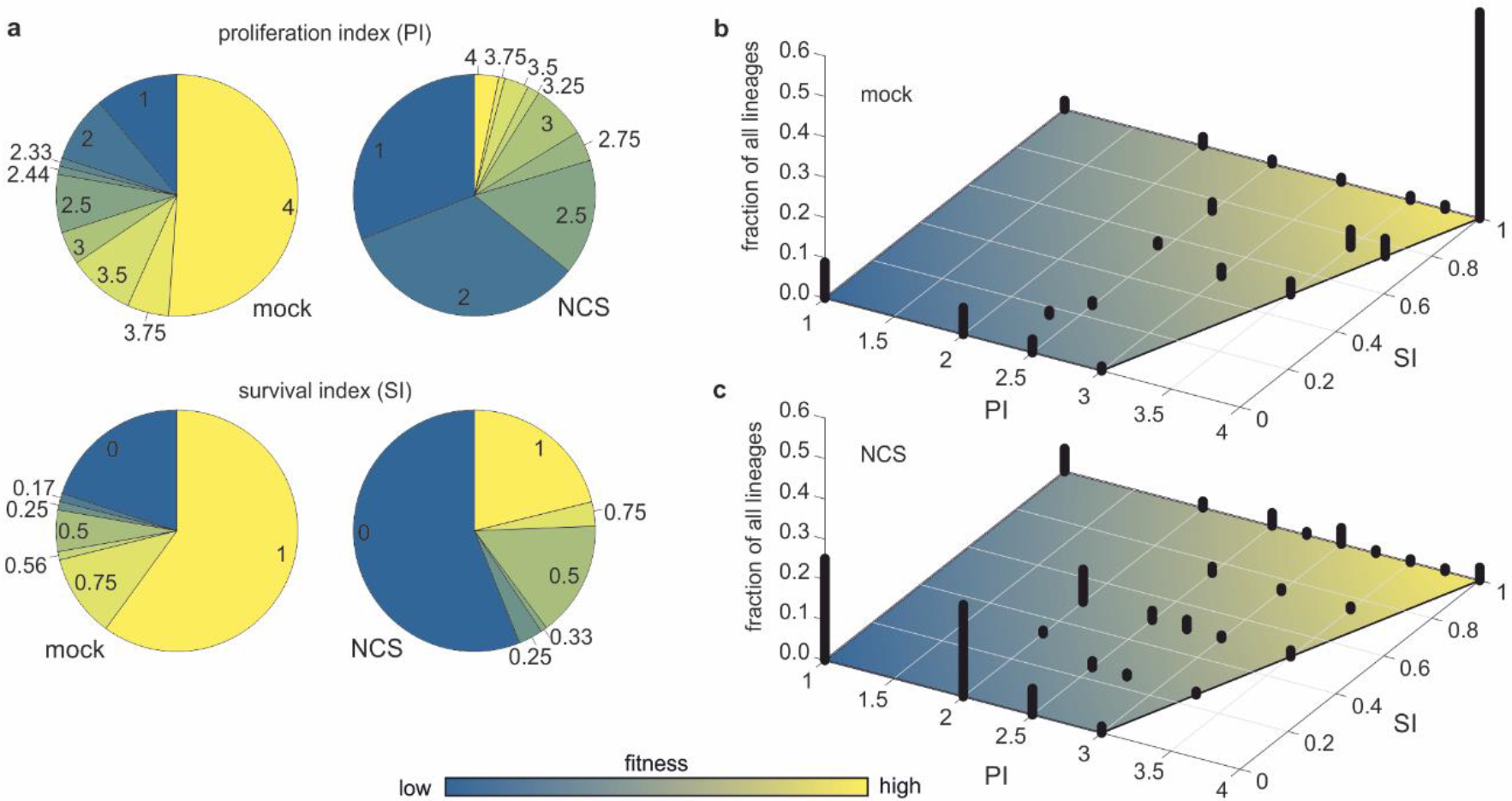
Non-genetic heterogeneity of cellular fitness after DNA damage. The fitness of a cell in response to DNA damage can be described by the capability of its progeny to proliferate and survive, quantitatively characterized by the proliferation (**a**, top) and survival indexes (**a**, bottom; see also Extended Data X for details). The prevalence of lineages in a lineage space depicting mock- (**b**) and NCS- (**c**) treated surviving and proliferating indexes exemplify the considerable heterogeneity of fitness that DNA damaged cells exhibit.

### The dangers and opportunities of cycling with damage

It has previously been argued that allowing DNA damage to pass through a checkpoint unrepaired is counterintuitively advantageous in a mutagenic environment allowing DNA repair and cell fate determination to be postponed to a more optimal time^18^. Indeed, our observation of delayed cell death may be such a deferred cell fate determination. However, although we have observed delayed cell death frequently drive lineage exhaustion there is a risk that each successful cell cycle will generate mutations that facilitate cancer evolution as cell cycle process act upon damaged DNA. Here, we modelled clonal dynamics featuring continuously high generational death rates comparable to those observed experimentally to examine how long and how many clones survive over time. Even at 55% death rates, ~20% of lineages can endure and even expand over two weeks of simulations, having undertaken many cell cycles in which further mutations can arise (**Fig. 5a-c**). Examining parental U2OS cells 5 days after NCS treatment for evidence of further genomic aberrations we identified the frequent production of micronuclei (**Fig. 5b**). Micronuclei are an aberrant genomic structure formed after broken DNA is incorrectly segregated during mitosis and are known to be involved in the generation of highly rearranged chromosomes via the process of chromothripsis^19,20^. Such processes occurring in tumours could therefore provide the substrate of mutations for tumour evolution leading to the emergence of more aggressive or resistant clones. On the other hand, these micronuclei were frequently positive for cGAS activation (**Fig. 5b**) indicating in accordance with recent studies^21,22^ that in a physiological environment such cells would provide an immunogenic stimulus and may encourage clearance by the immune system.

**Figure 5.**
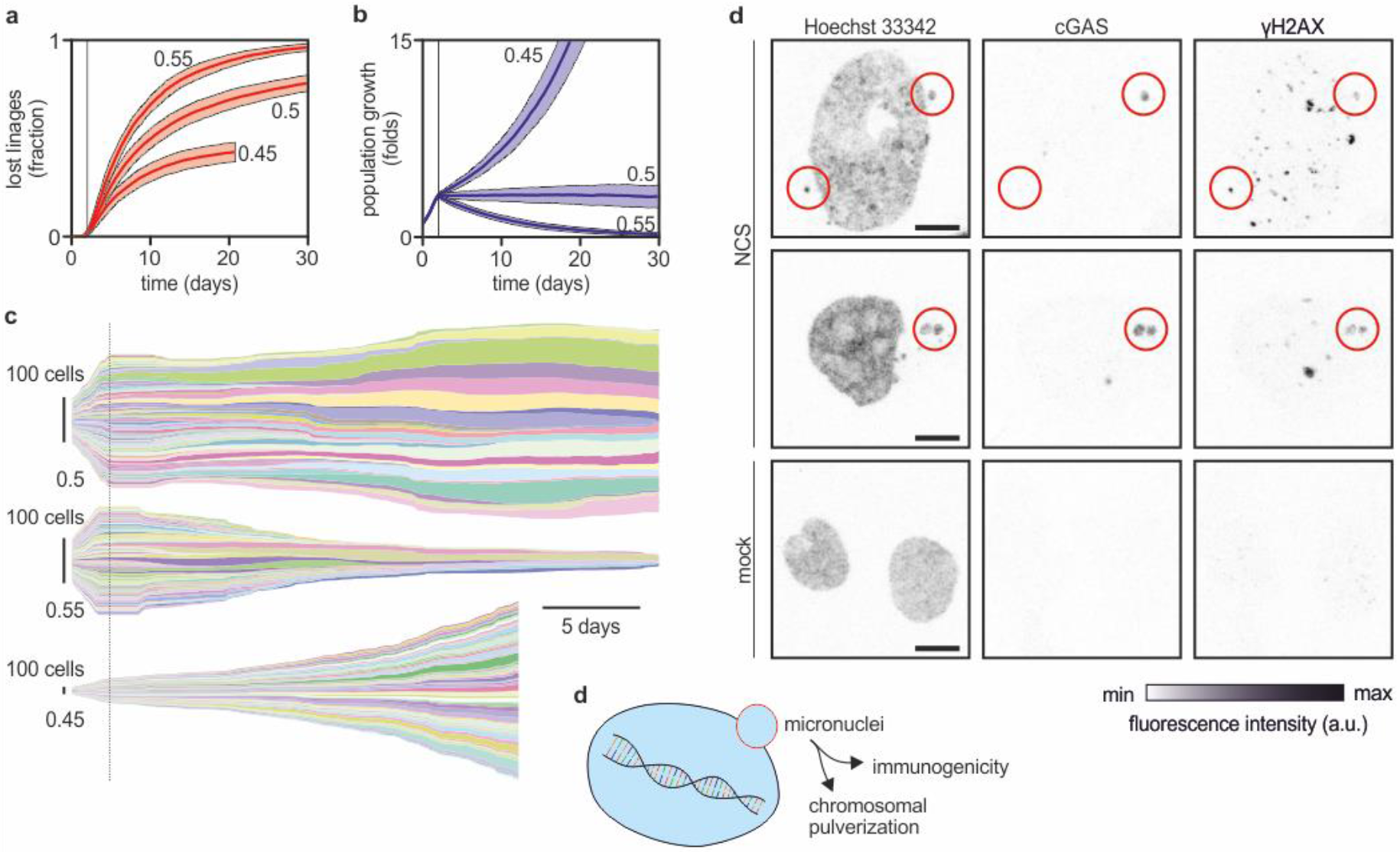
Dying later or growing stronger? Cancer cells that exhibit a decreased fidelity of the DNA damage checkpoint can cycle in the presence of DNA damage while shedding a fraction of their population (α) at every population as shown in clonal dynamics simulation (**a-c**). Panel **a**) and **b**) shows the fraction of exhausted lineages and the population fold increase, respectively, at different levels of cell death (45%, 50% and 55% per generation). Solid lines and shaded areas represent averages ± standard deviations computed over 100 repeats of Monte Carlo simulations. Panel **d**) depicts one example of clonal dynamics for each setting. Either if this process leads towards the exhaustion (α>0.5) or the expansion (α<0.5) of the cancer cell population, cycling with persistent DNA damage leads to micronuclei formation (**d**) and the activation of the cGAS pathway. Delayed cell death, therefore, provides both an opportunity for cancer cell to select resistant phenotypes through genomic rearrangements or an opportunity for clinical intervention (**d**).

## Discussion

Taken together, our results suggest that when non-ideal checkpoint fidelity permits DNA damaged cells to continue to divide, subsequent generations revisit cell fate decisions leading to delayed cell death, a phenomenon only poorly studied to date. Fidelity of checkpoints depends on various factors including the time at which damage occurred, the quantity and type of lesions, all factors that cannot be precisely controlled during clinical interventions. Moreover, fidelity of checkpoints depends also on non-genetic heterogeneity and it manifest itself in various forms, from failure to trigger a cell cycle arrest to variability in timing and negligence towards low level of damage or specific lesions. Eventually, non-ideal fidelity of checkpoints results into a vastly heterogeneous landscape of lineage fitness that can be revealed only through time-lapse lineage tracing of intact living cells and that it is commonly neglected.

Our results suggest persistent DNA damage could provide the signal that leads to the observed descendant cell death. However if so, the lack of cell cycle arrest we report in intermediate generations raises the interesting possibility that inherited DNA damage does not cause the canonical cell cycle checkpoint response perhaps via a yet to be identified checkpoint marking or memory process analogous to those demonstrated for replication stress^23–25^.

When treated cells do not commit to terminal cell fates but revisit cell fate decisions in later generations, we are challenged with risks in relation to clinical interventions, but also opportunities. High frequency of delayed cell death permits the extinction of many lineages correcting the initial failure of checkpoints to prevent cell cycling. However, cells cycling with DNA damage amplify genomic aberrations via the production of micronuclei from persistent DNA breaks. In a physiological context such mutations could provide the substrate for selection of more aggressive or resistant clones. However, the formation of micronuclei and resultant cGAS mediated immunogenic signal act as an ultimate tumour suppressor mechanism for DNA damaged descendant cell death, which may also be amenable to immunotherapeutic intervention.

## Material and Methods

### Cell culture

U2OS (ATCC®> HTB-96^™^) cells were obtained from ATCC and STR profiled by the Cambridge Institute CRUK core facilities. U2OS cells were grown in DMEM+GlutaMAX^™^-I supplemented with fetal bovine serum (FBS) (ratio 10:1 media to FBS) and Pen Strep. During microscopy U2OS cells were cultured in phenol-red free DMEM supplemented with FBS (ratio 10:1 media to FBS), sodium pyruvate (1mM Gibco 11360-070), Glutamax-I (Gibco. 35050-038) and Pen Strep or CO2-independent Leibowitz L15 medium supplemented with 10% FBS and Pen Strep. Cell cultures were maintained in Nunclon^™^ delta treated flasks, dishes and plates.

### Gene synthesis, plasmids and constructs

Gene synthesis was provided by ThermoFisher LifeTechnologies (GeneArt Gene synthesis and GeneArt Strings). mKO2-hCdt1(aa30-120) and mAG-hGeminin(aa1-110) gene sequences were synthesised according to the sequences detailed in (Sakaue-Sawano et al. 2008)^26^ (Genbank: AB370332.1 AB370333.1). mKO2 was substituted for mCherry and the resultant mCherry-hCdt1(aa30-120) was cloned downstream of the CMV promoter in the dual-promoter plasmid pBudCE4.1 mAG hGeminin(aa1-110) was cloned downstream of the Ef1α promoter. iRFP682 was synthesised according to the sequences detailed in (Shcherbakova & Verkhusha 2013) (Genbank: KC991143)^27^ and NLS tagged variants generated via restriction enzyme cloning into the plasmid backbone pcDNA3.1(-).

### Stable cell line generation

U2OS-Tracer cells were obtained via JetPrime^®^ transfection with pBudCE4.1 mCherry hCdt1(aa) mAG hGem(aa) and pcDNA3.1(−) (1+1)×NLS iRFP682 followed by selection with zeocin and G418. Monoclonal cell populations were obtained via fluorescence activated cell sorting which were then screened by microscopy for successful stable integration of the two plasmids.

### Neocarzinostatin treatments

Neocarzinostatin (NCS) was purchased from Sigma-Aldrich (Cas No. 9014-02-2, MDL No. MFCD01778130 Catalogue No. N9162) as a 0.5 mg/ml solution in 20 mM MES buffer (pH 5.5). NCS-containing media was prepared to the indicated concentrations while mock treatments were prepared with the equivalent volume of 20 mM MES buffer (pH 5.5). NCS concentrations were calibrated between different experimental assays to attempt to compare similar magnitudes of phenotype. This was necessary due to the difference between treating dishes with NCS and flowing volumes of NCS containing media over chamber slides. For example, concentrations of 100 ng/ml NCS applied to dishes appeared to be comparable to 30-45 ng/ml NCS applied to perfusion chamber slides in microscopy-based lineage tracing assays.

### Microscopy based lineage tracing - experimental setup

FUCCI microscopy lineage tracing experiments were carried out in Ibidi III 3D perfusion μ-slides and associated flow accessories from Ibidi. (Luer Lock Connector Female, Elbow Luer Connector, y tube fitting, Silicone Tubing. Catalogue numbers: 10825, 10802, 10827, 10841). U2OS-Tracer cells were seeded at a surface density of 3600 cells cm-2 and left to attach. The chamber slide was then charged with media. Approximately a day later flow tubing was attached, and fresh media washed through. Slides were then attached to the microscopy stage, fresh media passed through and imaging begun. One day after the start of image acquisition sufficient excess volumes of media, containing mock or NCS treatments (30/45 ng/ml NCS in experiments), to wash through the entire flow tubing and chamber slide system were passed through the tubing. An hour later similar excess volumes of media were washed through the tubing to remove any residual NCS containing media. After a number of pilot experiments, we noticed that the magnitude of phenotype induced by NCS treatments was variable between different experiments, although always present. Therefore, aiming to standardize our experimental conditions, as batch-to-batch effectiveness of NCS and perfusion might change, we countered this effect by performing a titration within each experiment and selecting those concentrations which displayed a moderate phenotype, sufficient to induce a cell cycle arrest and cell death but not so severe that no cells die.

### Image acquisition

Widefield microscopy was carried out using a Nikon Eclipse Ti inverted microscope controlled by NIS-Elements AR software (Version 4.30.02). Imaging was achieved with a Zyla sCMOS camera (Andor Instruments). Images were acquired at 30 minutes intervals with 20x air objective lens magnification, 2560×2160 pixel size. Four channels were acquired a bright field transmitted light image and three fluorescence images, far-red (Cy5 filterset), green (FITC filterset) and red fluorescence (TRITC filterset). Time-lapse microscopy continued for 6 days following the start of the experiment.

### Image segmentation

Fluorescence images, using either just the nuclear marker or the nuclear marker and the G1 and SG2 FUCCI probes, were segmented using the NIS-Elements AR built in segmentation function followed by morphological opening using the command: “OpenBinaryND(8,3,1,1,1);” and separation of objects using: “MorphoSeparateObjectsND(15,7,1,1,1);”

The resultant image overlays were exported as tif files and nuclear masks extracted from these. Where three fluorescent channels were used independently for segmentation the resultant three masks were combined to create a single final mask.

### LinSense

The generation and curation of lineages was performed using LinSense, a custom MATLAB software we developed and that is freely available from the GitHub repository *alesposito/LinSense*.

Briefly, the nuclear masks produced from image segmentation were tracked through time. The centroids of each nucleus identified were compared to previous images and tracking achieved by comparing the distances between them. The resultant tracked nuclei were used to make measurements of average FUCCI probe fluorescence for each cell. The resultant fluorescence traces were then corrected by manual validation to account for erroneous segmentation and tracking. Cell death and division were assessed by eye and their timings and identities of daughter cells recorded. Cell lineages were analysed to the third generation, the Granddaughter cells, following treatment.

The G1/S phase transition was estimated based on the peaking of the red fluorescent G1 probe mCherry-hCdt1(aa30-120). This was done by assessing both the raw fluorescence measurements and a median filtered version of the data that smoothed experimental noise. Multiple apparent G1 phases could be determined in cells where green SG2 probe fluorescence declined and red G1 probe fluorescence increased without the cell having gone through mitosis.

### Generational tracing by flow cytometry

A sufficient number of cells for each experiment were pelleted and resuspended in a 37°C 10 μM CellTrace^™^ Yellow (ThermoFisher Scientific Catalogue No. C34567) solution in PBS at a concentration not greater than one million cells per ml and incubated at 37°C for 20 minutes. Residual dye was quenched by the addition of 5 volumes of culture media and cells incubated for a further 5 minutes at 37°C. Cells were pelleted, resuspended in complete media, and seeded and seeded in Nunclon Delta surface 6cm dishes at a surface density of typically 9000 cells cm-2 (approximately 190,000 cells) in a total volume of 4mls of complete culture medium. Cells were left for approximately 24 hours before treatment to allow attachment and recovery from seeding. Cells were trypsinised and to produce a population of cells in suspended.

### NCS treatment

At the start of the experiment cell culture media was aspirated from all dishes apart from the 0 hour samples and replaced treated with 3ml of culture medium containing the desired concentration of NCS or an equivalent volume of 20mM MES buffer (pH 5.5) in the case of control experiments. 1-2 hours later this was removed, dishes were washed with PBS and 4mls of fresh culture medium added. Any residual unreacted and undegraded NCS should therefore have been reduced to negligible levels.

Dishes were seeded allowing samples to be taken at 24-hour intervals for each condition. U2OS experiments were conducted with triplicate dishes for each timepoint and condition. Additional dishes were seeded to allow collection of the cells bound to the dish for cell cycle profiling. During sample harvesting cells were trypsinised collecting their culture media and PBS washes. Trypsinised cells were then washed off the dish, collected together with culture media and PBS washes and pelleted. Cells were then resuspended in PBS, a fraction of which was taken to be counted, the remainder mixed with formaldehyde in PBS to a final concentration of 2% formaldehyde and incubated for 15 minutes on ice. Cells were then washed twice in ice-cold PBS and stored at 4°C in PBS until analysis.

In some experiments cell cycle profiles were obtained from separately harvested cells which were fixed in 70% EtOH in water and stored at −20°C until required after which they were washed once in PBS, stained with Hoechst 33342 (Sigma-Aldrich Catalogue No. H3570) and analysed.

The fraction of the cell samples dedicated to population counting was diluted in PBS or Leibowitz L15 medium supplemented with FBS and Pen Strep, and analysed on a Beckman Coulter “Vi-cell-XR cell viability analyzer” machine using default cell parameters. This machine provides information both on the estimated total cell concentration and the viable cell concentration based on trypan blue based cell viability discrimination. Both measures were subsequently used for analysing the relative changes in population over time.

Flow cytometry events were acquired as above. Following gating, fluorescent curves representing the CellTrace stained cells were obtained for each sample and the median value extracted to permit fluorescence comparison across samples. Fluorescence measurements were paired with their corresponding population estimates and plotted as indicated in the results. Linear regression was carried out by determining the line of best fit constrained by passing through the origin, i.e. the first time point measurements. Where experiments were carried out with multiple replicates per time point the origin was defined by the average of the first time point replicates.

### Clonal dynamics simulations

A simple simulation of clonal dynamics was utilized to illustrate the consequences of cell proliferation with delayed cell death on cell lineages. The software is available as the GitHub repository *alesposito/CloE-DDR*. Briefly, we first generate 100 virtual cells with a randomized cell cycle. Cell cycle lengths are drawn from a gaussian distribution modelled on the U2OS cells with 24.5 hrs average and 5 hrs standard deviation. Cells are then left to divide with randomized lengths at any cycle for thirty days with a time resolution of one hour. After two days during which cells were simulated with a probability of cell death equal to zero, the probability of cell death per cell cycle was set to arbitrary values, here 0.45, 0.5 and 0.55. CloE-DDR is also creating lineage trees compatible to LinSense analysis for future applications. Here, we show only clone size as a function of time. Simulations exceeding 2500 lineages were interrupted to keep CloE-DDR compatible with standard computers. For each parameter set, the Monte Carlo simulations were repeated 100 times. Here, we show the average and standard deviation of relevant statistics computed over the 100 repeats, such as the fold increase of population (normalized to the initial 100 cells) and the fraction of lineages that is exhausted.

### Micronuclei assay

Cells were seeded in Nunclon Delta surface 6 cm dishes on glass coverslips at a surface density of typically 9000 cells cm-2 (approximately 190,000 cells) in a total volume of 4mls of complete culture medium. Cells were left for approximately 24 hours before treatment to allow attachment and recovery from seeding. At the start of the experiment cell culture media was aspirated from all dishes and replaced with 3ml of culture medium containing the desired concentration of NCS or an equivalent volume of 20mM MES buffer (pH 5.5) in the case of control experiments. 1-2 hours later this was removed, dishes were washed with PBS and 4mls of fresh culture medium added. Any residual unreacted and undegraded NCS should therefore have been reduced to negligible levels. Samples were collected two hours and five days after NCS treatment and fixed with 4% formaldehyde in PBS at room temperature for 10-15 minutes. Cells were then washed twice in PBS and stored at 4°C in PBS until developed to detect γH2AX and cGAS by immunofluorescence (see below). Coverslips were also stained Hoechst 33342 (Sigma-Aldrich Catalogue No. H3570) for DNA detection. Coverslips were analysed on a Leica SP5 microscope. Images were acquired with a photo-multiplier tube detector using a 40× objective lens and the settings 512×512 pixel image size, 400 Hz scan speed, 2× zoom, 2-4× line or frame averaging with sequential scanning. Laser power was adjusted based on the appropriate level for sample brightness, permitting clear visualisation of fluorophores but avoiding pixel saturation.

### Immunofluorescence

Cells were fixed using a solution of 4% paraformaldehyde, permeabilised (0.1% TritonX-100 in PBS 5 mins at room temperature), blocked (Blocking solution: 2% BSA, 0.2% Tween, 0.1% Triton X-100 in PBS 60-90 mins at room temperature) and incubated with primary antibodies in blocking solution for 1 hour at 37°C (anti-γH2AX (Millipore JBW301) 1:2000). Cells were washed thrice (washing solution: 0.2% Tween, 0.1% Triton X-100 in PBS) and incubated with secondary antibody and Hoechst dye in blocking solution (Hoechst 33342 1:1000, Alexafluor 594 goat anti-mouse IgG 1:500), washed in washing solution twice, washed twice in PBS, left in PBS and analysed by microscopy.

cGAS staining was carried out as described in Mackenzie *et al.*^21^. This protocol was adapted by the additional inclusion of anti-γH2AX (Millipore JBW301) at a 1:2000 dilution.

## Acknowledgments

C.J.C was supported by a CRUK Cambridge Cancer Centre studentship, A.E., by an MRC core award, to A.R.V. We would like to thank John Saganty for help on data analysis related to this project.

